# 2D Label-free Prediction of Multiple Organelles Across Different Transmitted-light Microscopy Images with Bag-of-Experts

**DOI:** 10.1101/2024.05.24.595656

**Authors:** Yu Zhou, Shuo Zhao, Justin Sonneck, Jianxu Chen

## Abstract

Label-free prediction has emerged as a significant application of artificial intelligence (AI) in the field of bioimaging, which aims to predict the localization of specific organelles directly from readily-accessible transmitted-light images, thereby alleviating the need for acquiring fluorescent images. Despite the existence of numerous research, in practice, the high variability in imaging conditions, modalities, and resolutions poses a challenge to the final prediction. In this study, we propose a “Bag-of-Experts” strategy, targeting at different organelles, with self-supervised pre-training. The comprehensive experimentation showcases that our model is agnostic to the transmitted-light image modalities and the imaging conditions, to certain extent, indicating considerable generalizability. The code is released at: https://github.com/MMV-Lab/LightMyCells

## 1. INTRODUCTION

In the past decade, AI has revolutionized the bioimaging field from many directions, e.g., image classification [1], image segmentation [2] and image restoration [3]. One important application is to directly predict the localization of specific organelles from transmitted-light (TL) microscopy images. Comparing to fluorescence microscopy, the TL images does not require fluorescent labeling and therefore are low-cost and can greatly reduce the difficulty in sample preparation and the phototoxicity during image acquisition. Virtually predicting organelle localization also breaks the practical limitation of the maximum number of organelles that can be simultaneously visualized by fluorescence proteins, and therefore enables a major step toward understanding the cell as an integrated system. The proof-of-concept of this technique first appeared in two works [4, 5], and therefore two terms, “label-free prediction” [4] and “in-silico labeling” [5], are interchangeably used in the bioimaging field. We use “label-free prediction” to refer this technique in this paper.

Even though the great potential of the label-free prediction method, its practical adoption for making integrated biological discoveries is still limited, mostly due to the unsatisfactory prediction accuracy. As shown in the recent work [6], the evaluation is much more than conventional image reconstruction metrics, like SSIM, and has to be application-appropriate [7]. In the bioimaging community, there is still no public benchmark or well-established evaluation methodologies for the label-free technique, yet. The “Light My Cells” challenge[8] offers a step-stone towards this direction by establishing a set of 2D benchmark datasets with large variations (e.g., different types of TL images, different instruments, etc.). In the paper, we will present the method we used in this challenge, summarize our results qualitatively and quantitatively, and at the end discuss limitations and outlooks.

## 2. DATA DESCRIPTION

The full details of the dataset can be found on the official challenge website[8]. Here, we provide a brief summary. There are about 57,000 2D images in total (with 4600 fluorescent images as ground truth), with 95% for training and 5% held-out for final evaluation. The images were acquired from different biological studies, and therefore may be of different magnification, from different instruments or even different acquisition sites. The dataset is organized as a series of TL image and fluorescent image pairs. The TL images could be bright field, phase contrast or differential interference contrast (DIC), while the corresponding fluorescent images could be of one of four different organelles, nucleus, mitochondria, tubulin, and actin. The TL images may come as a single Z-slice (may or may not be of best Z-focus) from a Z-stack, but the fluorescent images are always 2D slice of best Z-focus (as determined by [9]), aiming to evaluate the generalizability and sensitivity of the label-free prediction models.

## 3. METHODOLOGY

The key idea of our method is to develop a single “Bag-of-Experts” model to predict from any given TL image, regardless of bright field, phase contrast, or DIC into four different images, each of a different organelle. The basic idea is to find the “expert” sub-model for each label-free prediction task. The development consists of three main stages: data cleaning, training individual experts, and integrating bag-of-experts, as elaborated below.

### 3.1. Data Cleaning

Data cleaning is necessary, considering the size and diversity of the training set. We implemented a semi-automatic data cleaning strategy. First, we employed the CleanVision tool^1^ as an automatic quality control (QC) step to detect potential data issues within the dataset, such as low information (low entropy in pixel values), duplication or blurry. Following QC, we arranged the ground truth images based on their gradient magnitude. We then conducted manual inspection of samples with the lowest scores, as lower values typically indicate fewer information content, until no further issues were identified. The image pairs with bad ground truth were discarded. In total, 77.23% images within the dataset were remained.

### 3.2. Training Individual Experts

Given four different organelles in the dataset (nucleus, mitochondria, tubulin, and actin), we created four sub-tasks, in each of which we conduct a sequence of experiments on different types of models (e.g., basic convolutional neural networks in MMV Im2Im package [10], transformer models, etc.), different pre-training strategies (e.g., training from scratch, pre-training using self-supervised training strategies like MAE [11]), different learning strategies (e.g., different optimizers and different learning hyper-parameters), etc. This resulted in a collection of experts for each sub-task. We then picked up the expert sub-model for each sub-task based on the performance on the evaluation dataset.

### 3.3. Integrating Bag-of-Experts

The final model was a bag of experts. Each expert was responsible for a specific organelle type, and was selected from all candidate experts for the corresponding sub-task. During inference, the meta-data from the input image can trigger the responsible experts.

## 4. EXPERIMENTS

### 4.1. Implementation details

#### 4.1.1. Model Specification

There are mainly two types of experts in our Bag-of-Experts model, namely UNet [12] and UNETR [13]. The former is the classic CNN-based methods, while the latter one has a similar architecture, but substituting the encoder with a ViT [14] backbone for the advancing feature extraction capability. For each expert, we also tuned with different architecture parameters to achieve optimal performance.

#### 4.1.2. Data Processing

We employed two different data processing pipeline for CNN-based methods and transformer-based methods. For CNN-based methods, the input image was firstly normalized to X ∼ 𝒩 (0, 1) and then randomly cropped and sampled as a list containing 8 patches with the size of 512 × 512. Some data augmentation methods were also adopted during training time, e.g, random flipping. We applied a sliding-window strategy when applying on the large-size input images, with the overlap ratio between the adjacent patches as 0.2.

For transformer-based methods, we transformed the data to RGB format and normalized the input data with the statistics from MAE. Then the image was randomly cropped and resized to 512 × 512. The same augmentation method was also applied. Noted that the image will not be cropped if the original size is less than 512. During the inference time, considering the global attention nature of ViT, we just interpolated the input image to 512× 512 and resized the prediction back to the original size, instead of applying the sliding-window strategy. We hypothesize that interpolating TL images, due to their sparse semantic information, will not incur significant information loss while significantly accelerating training and reducing memory usage.

#### 4.1.3. Pre-training Strategies

Substantial amount of research demonstrated that proper pretraining would enhance the performance of the downstream tasks [11, 15]. So in this study, we pre-trained the ViT-based backbone of the UNETR model using MAE [11], a well-known self-supervised pre-training strategy, in order to better exploit the semantic information of input TL images. A typical MAE network consists of a ViT-based encoder and a simple decoder. We included all the TL images as our training set and masked out 75% of the input image randomly using 16 × 16 small patches. We followed the MAE training pipeline to reconstruct the original image based on the remaining pixels. At the end, the pre-trained encoder could be used as the feature-extraction backbone for downstream label-free tasks.

#### 4.1.4. Learning Strategies

For CNN-based methods, we trained the model from scratch and used SmoothL1Loss as the criterion. Adam optimizer was chose with a 0.001 initial learning rate. The maximum epoch was 5000 with early stopping for avoiding over-fitting.

Regarding the transformer-based methods, the model was pre-trained and fine-tuned with 1000 epochs. The lightweight Lion optimizer [16] was chosen combined with the cosine annealing scheduler. We set the initial learning rate 70% less than what we used in the methods with Adam optimizer, as suggested in [16]. The details are listed in table 1. Note that the hyperparameter configuration remains consistent across all sub-models sharing the same model structure.

**Table 1.** Characteristics of the Learning Setups.

| Learning Setups | UNet | UNETR |
| --- | --- | --- |
| Initialization | He normal init | pretrained (encoder)<br>+<br>He normal init (decoder) |
| Batch size | $8 \times 8$ (patches) | 16 or 24 |
| Total epochs | 5000 | 1000 |
| Loss function | SmoothL1 | SmoothL1 |
| Optimizer | Adam | Lion |
| Scheduler | ExponentialLR<br>(gamma = 0.98) | CosineAnnealing<br>($T_{mul} = 2$, decay = 0.9) |
| Initial learning rate (lr) | 1e-3 | 3e-4 |
| Implementation | mmv_im2im | PyTorch + Fabric <sup>2</sup> |
| Precision | fp32 | bf16 |

## 5. RESULTS

Our bag-of-experts strategy is evaluated from five aspects, namely Mean Absolute Error (MAE), Structural similarity index measure (SSIM), Pearson correlation coefficient (PCC), Euclidean distance (ED), Cosine distance (CD). The first three metrics focus on the overall similarity while the last two highlight texture features, which are usually used for traditional cellular phenotyping. We take all the slices into account instead of 0-5 deviations of the focus plane, since we have no information of the focus location.

For all four types of organelles, the predictions generated by the pretrained UNETR model consistently outperformed those of the classic UNet model trained from scratch. This superiority can be attributed to the incorporation of the ViT encoder and the MAE-based pretraining methodology, as discussed in the subsequent section. In terms of SSIM, the mitochondria prediction reach the highest (0.8913), while the nucleus prediction achieving the best PCC value (0.9676). Overall, the predictions were quite similar to the corresponding fluorescent images.

## 6. DISCUSSION

The generalization ability of our strategy is displayed in fig. 1. The individual experts were trained using both the whole dataset and the individual image modality (BF, DIC, PC), respectively. In terms of SSIM, the model trained by all the input images have equivalent or slightly worse performance across all organelles and TL modalities. The metric in the prediction from BF/PC to tubulin even outperform the separately trained model. Besides, all the models were trained based on all 30 study cohorts. This showcases the robustness of our model to different imaging cohorts and modalities.

**Fig. 1.**
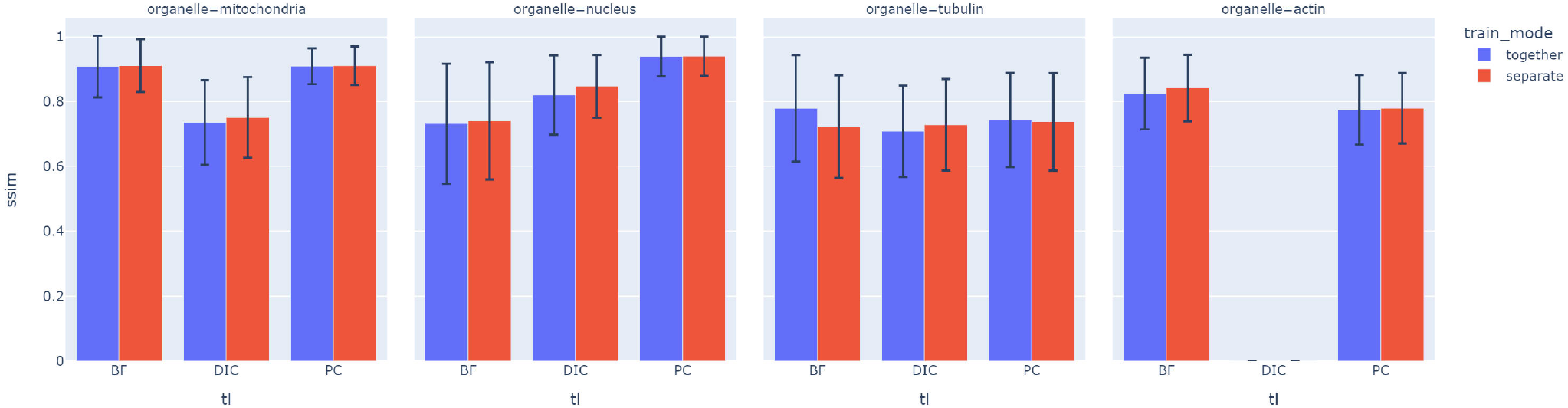
SSIM comparison between different training modes.

We also did some ablation study to evaluate the effectiveness of the MAE pre-training, as illustrated in table 2. For tubulin, the accuracy will drop heavily in all the input image modalities for all the metrics, if no MAE pre-training is applied beforehand. This means MAE pre-training can grasp the semantic information across different image modalities and contribute largely to the generalization ability of the model.

**Table 2.** Performance comparison: with and w/o MAE pre-training for tubulin prediction.

| Modality | Pretrain | SSIM | PCC |
| --- | --- | --- | --- |
| BF | ✓ | **0.7744** ± 0.1541 | **0.9602** ± 0.0145 |
| BF | ✗ | 0.7050 ± 0.1555 | 0.8669 ± 0.0720 |
| DIC | ✓ | **0.7354** ± 0.1359 | **0.9669** ± 0.0336 |
| DIC | ✗ | 0.5653 ± 0.1247 | 0.7423 ± 0.2400 |
| PC | ✓ | **0.7550** ± 0.1369 | **0.9567** ± 0.0148 |
| PC | ✗ | 0.6571 ± 0.1409 | 0.8551 ± 0.0690 |
| overall | ✓ | **0.7447** ± 0.1387 | **0.9633** ± 0.0283 |
| overall | ✗ | 0.6049 ± 0.1426 | 0.7864 ± 0.2018 |

**Table 3.** Model performance on label-free prediction of different organelle types.

| Organelle | Submodels | MAE | SSIM | PCC | ED | CD |
| --- | --- | --- | --- | --- | --- | --- |
| Mitochondria | UNETR | $0.0200 \pm 0.0182$ | $0.8913 \pm 0.1043$ | $0.9625 \pm 0.0619$ | $50.9322 \pm 63.2961$ | $0.1342 \pm 0.1599$ |
| Nucleus | UNETR | $0.0238 \pm 0.0189$ | $0.7818 \pm 0.1818$ | $0.9676 \pm 0.0392$ | $48.3074 \pm 40.7744$ | $0.1360 \pm 0.1663$ |
| Tubulin | UNETR | \ | $0.7447 \pm 0.1387$ | $0.9633 \pm 0.0283$ | \ | \ |
| Actin | UNETR | \ | $0.7889 \pm 0.1105$ | $0.9287 \pm 0.1635$ | \ | \ |

Our method indeed has some limitations. For example, we cannot predict all four kinds of organelle localization from only one expert sub-model, which will be in our future research plan. Meanwhile, regarding the variability of input images, we do not handle the class imbalance, which may result in worse prediction in the images of minority classes.

## 7. CONCLUSION

This study investigated the difficulty in the label-free prediction task due to the high variability of the transmitted-light images and proposed a robust and generalizable algorithm, which is agnostic to the imaging cohort and the input modalities. We applied a “Bag-of-Experts” strategy to target specifically at different organelle types. To overcome the heterogeneity of the input images, MAE pre-training technique was also employed to better extract the semantic information of the whole dataset. The result shows that our prediction is plausible and stable across different image modalities and study cohorts.

## 8. COMPLIANCE WITH ETHICAL STANDARDS

The authors declare no competing interests.

## 9. ACKNOWLEDGMENTS

All authors are funded by the Federal Ministry of Education and Research (BMBF) in Germany under the funding reference 161L0272, and also supported by the Ministry of Culture and Science (MKW) of the State of North Rhine-Westphalia.

## Footnotes

1 https://github.com/cleanlab/cleanvision

